# Bacterial DNA induces the formation of heat-resistant disease-associated “Tetz-proteins” in human plasma

**DOI:** 10.1101/604090

**Authors:** Victor Tetz, George V. Tetz

## Abstract

Our study demonstrated for the first time that bacterial extracellular DNA (eDNA) can change the thermal behaviour of specific human plasma proteins, leading to an elevation of the heat-resistant protein fraction, as well as to *de novo* acquisition of heat-resistance. In fact, the majority of these proteins were not known to be heat-resistant and nor do they possess any prion-like domain. Proteins found to become heat-resistant following DNA exposure were named “Tetz-proteins”.

Interestingly, plasma proteins that become heat-resistant following treatment with bacterial eDNA are known to be associated with cancer progression. Therefore, we analysed the heat-resistant proteome in the plasma of healthy subjects and in patients with pancreatic cancer and found that exposure to bacterial eDNA made the proteome of healthy subjects more similar to that of cancer patients. These findings open a discussion on the possible novel role of eDNA in disease development following its interaction with specific proteins, including those involved in multifactorial diseases such as cancer.

## Introduction

The role of microbiota is being revisited due to its emerging role in pathologies that were previously considered non-microbial [1,2]. For instance, bacteriophages have been recently found to be associated with the development of specific human diseases, such as Parkinson’s disease and type 1 diabetes [3-5]. Particular attention to diseases triggering has also been drawn by pathogen-associated molecular patterns (PAMPs), mainly represented by components of microbial biofilms, including those of the gut microbiota [6]. One example is bacterial extracellular DNA (eDNA). Bacteria produce large amounts of eDNA that plays a multifunctional role in microbial biofilms, as a structural component, a nutrient during starvation, a promoter of colony spreading, and a pool for horizontal gene transfer [7-9]. eDNA is also known to affect bacterial protein modification in biofilm matrix, as exemplified by its role in the conversion of bacterial extracellular polymeric amyloids into β-sheet-rich structures with prion-like characteristics, such as self-propagation and resistance to proteases and heat [10-12]. Heat resistance is a hallmark of prion proteins, although its biological significance is not clear. Notably, heat resistance is not exclusive of prion proteins and also characterizes mammalian proteins that are not implicated in heat-shock events, which makes this phenomenon even more puzzling [13,14].

The properties of bacterial eDNA have been poorly investigated, except for its actions in the context of microbial biofilms. On the other hand, the chances that the eDNA secreted from microbial communities interacts with human proteins are relatively high. For example, eDNA released during biofilm spreading or lytic bacteriophage infection can enter systemic circulation by different pathways and is facilitated by the altered intestinal permeability that is accompanied by the increased absorption of PAMPs [15-17]. Increasing evidence shows that impaired gut barrier dysfunction is an important determinant for the increase in circulating bacterial DNA that is associated with different diseases. Indeed, increased levels of both bacterial eDNA and human cfDNA characterize various pathological human conditions including cancer, stroke, traumas, autoimmune disorders, and sepsis [18-21].

Another way by which PAMPs can enter biological fluids is their release from bacteria localized within the “internal environment” such as brain or placenta [22-25]. Moreover, DNA can be released inside eukaryotic cells from obligate and facultative intracellular bacteria [26, 27].

Thus, despite the fact that interactions between bacterial eDNA and humans are very likely to occur, the effects of bacterial eDNA within body fluids are poorly studied, except for the CpG motif-induced activation of proinflammatory reactions through Toll-like receptor 9 [28]. In this study, we evaluated a novel effect of bacterial eDNA on blood plasma proteins, which resulted in the alteration of the heat resistance of these proteins.

## Results

### eDNA-induced alteration of protein heat resistance in the plasma of healthy controls

We first studied the effects of DNA on the thermal behaviour of proteins from the plasma of healthy individuals. Most proteins were aggregated after boiling, and the supernatant contained heat-resistant fractions of over 100 proteins. The identified heat-resistant proteins had a molecular weight in a range of 8 kDa to 263 kDa. Treatment with bacterial and human buffy coat DNA altered the composition of the heat-resistant protein fraction. We first verified the levels of which plasma proteins, identified as heat-resistant before the treatment with DNA, were increased following DNA exposure in at least one healthy control (Table 1).

**Table 1.** Heat-resistant proteins of healthy controls whose amount increased following treatment with different DNAs*.

| N | Accession No<br>UniProt | Uniprot Accession | Protein name |
| --- | --- | --- | --- |
| **eDNA of *P. aeruginosa*** |  |  |  |
| 1 | P02768 | ALBU_HUMAN | Serum albumin |
| 2 | P02751 | FINC_HUMAN | Fibronectin |
| 3 | B4E1Z4 | B4E1Z4_HUMA | cDNA FLJ55673, highly similar to Complement factor B |
| 4 | P02774 | VTDB_HUMAN | Vitamin D-binding protein |
| 5 | P01859 | IGHG2_HUMAN | Immunoglobulin heavy constant gamma 2 |
| 6 | P00747 | PLMN_HUMAN | Plasminogen |
| 8 | Q14624 | ITIH4_HUMAN | Inter-alpha-trypsin inhibitor heavy chain H4 |
| 9 | Q5T987 | ITIH2_HUMAN | Inter-alpha-trypsin inhibitor heavy chain H2 |
| 12 | P04114 | APOB_HUMAN | Apolipoprotein B-100 |
| 13 | O14791 | APOL1_HUMAN | Apolipoprotein L1 |
| 15 | P19652 | A1AG2_HUMAN | Alpha-1-acid glycoprotein 2 |
| 16 | P20851 | C4BPB_HUMAN | C4b-binding protein beta chain |
| 3 | P01857 | IGHG1_HUMAN | Immunoglobulin heavy constant gamma 1 |
| **eDNA of *S.aureus*** |  |  |  |
| 17 | P02652 | APOA2_HUMAN | Apolipoprotein A-II |
| **eDNA of *E.coli*** |  |  |  |
| 18 | P19652 | A1AG2_HUMAN | Alpha-1-acid glycoprotein 2 |
| 19 | P04114 | APOB_HUMAN | Apolipoprotein B-100 |
| 20 | P20851 | C4BPB_HUMAN | C4b-binding protein beta chain |

\*Significant fold change in the level of heat-resistant proteins between normal plasma and plasma treated with eDNA for the proteins with spectrum counts <200 and over 30% increase for the proteins with spectrum counts $\geq 200$ .\*

We next measured the increase in heat-resistant protein fractions following the treatment of plasma with bacterial eDNA. The highest increase in heat-resistant fractions of different unrelated proteins was registered after the incubation with eDNA of *Pseudomonas aeruginosa*. Notably, eDNA from different bacteria produced distinct effects. Indeed, the exposure to eDNA from *Staphylococcus aureus* resulted in a selective increase in heat-resistant APOA2, which was not observed after treatment with eDNA from other bacteria. Under the same conditions, *E. coli* eDNA increased the heat-resistant fractions of A1AG2, APOB, and C4BP; however, the latter heat-resistant fractions were also increased after exposure to *P. aeruginosa* eDNA.

Intriguingly, specific proteins that did not exhibit a heat-resistant fraction in untreated plasma samples became heat-resistant following eDNA exposure. Table 2 lists the proteins that displayed such a behaviour in at least one of the plasma samples.

**Table 2.** Proteins that became heat-resistant following eDNA treatment but had no heat resistant fractions before.

| N | Accession No<br>UniProt | Uniprot Accession | Protein name |
| --- | --- | --- | --- |
| **eDNA of *P.aeruginosa*** |  |  |  |
| 1 | P69905 | HBA_HUMAN | Hemoglobin subunit alpha |
| 2 | Q03591 | FHR1_HUMAN | Complement factor H-related protein 1 |
| 3 | P01031 | CO5_HUMAN | Complement C5 |
| 4 | A0M8Q6 | IGLC7_HUMAN | Immunoglobulin lambda constant 7 |
| 5 | O43866 | CD5L_HUMAN | CD5 antigen-like |
| 6 | P49908 | SEPP1_HUMAN | Selenoprotein P |
| 7 | P0DOY3 | IGLC3_HUMAN | Immunoglobulin lambda constant 3 |
| 8 | P63241 | IF5A1_HUMAN | Eukaryotic translation initiation factor 5A-1 |
| 9 | P04264 | K2C1_HUMAN | Cluster of Keratin, type II cytoskeletal 1 |
| 10 | P35527 | K1C9_HUMAN | Keratin, type I cytoskeletal 9 |
| 11 | P13645 | K1C10_HUMAN | Keratin, type I cytoskeletal 10 |
| 12 | A0A075B6S5 | KV127_HUMAN | Immunoglobulin kappa variable 1-27 |
| **eDNA of E.coli** |  |  |  |
| 1 | Q9P2D1 | CHD7_HUMAN | Chromodomain-helicase-DNA-binding protein 7 |
| 2 | Q9UGM5 | FETUB_HUMAN | Fetuin-B |
| 3 | P01857 | IGHG1_HUMAN | Immunoglobulin heavy constant gamma 1 |
| 4 | P01861 | IGHG4_HUMAN | Immunoglobulin heavy constant gamma 4 |
| 5 | P01718 | IGLV3-27 | Immunoglobulin lambda variable 3-27 |
| 6 | P20151 | KLK2 | Kallikrein-2 |
| 7 | Q8TBK2 | SETD6_HUMAN | N-lysine methyltransferase SETD6 |
| 8 | P18583 | SON_HUMAN | Protein SON |
| 9 | O95980 | RECK_HUMAN | Reversion-inducing cysteine-rich protein with Kazal motifs |
| 10 | P02787 | TRFE_HUMAN | Serotransferrin |
| 11 | P49908 | SEPP1_HUMAN | Selenoprotein P |
| 12 | P0DOY3 | IGLC3_HUMAN | Immunoglobulin lambda constant 3 |
| 13 | P63241 | IF5A1_HUMAN | Eukaryotic translation initiation factor 5A-1 |
| 14 | P13645 | K1C10_HUMAN | Keratin, type I cytoskeletal 10 |
| **Human DNA** |  |  |  |
| 1 | P04264 | K2C1_HUMAN | Cluster of Keratin, type II cytoskeletal 1 |
| 2 | P35527 | K1C9_HUMAN | Keratin, type I cytoskeletal 9 |
| 3 | P13645 | K1C10_HUMAN | Keratin, type I cytoskeletal 10 |

These findings clearly demonstrated that human DNA and eDNA from different bacteria had a distinct influence on the generation of heat-resistant protein fractions.

To further analyse the correlation between DNA exposure and acquisition of heat resistance, we constructed a heat map summarizing the impact of different DNAs on the thermal behaviour of proteins (Fig 1)

**Figure 1.**
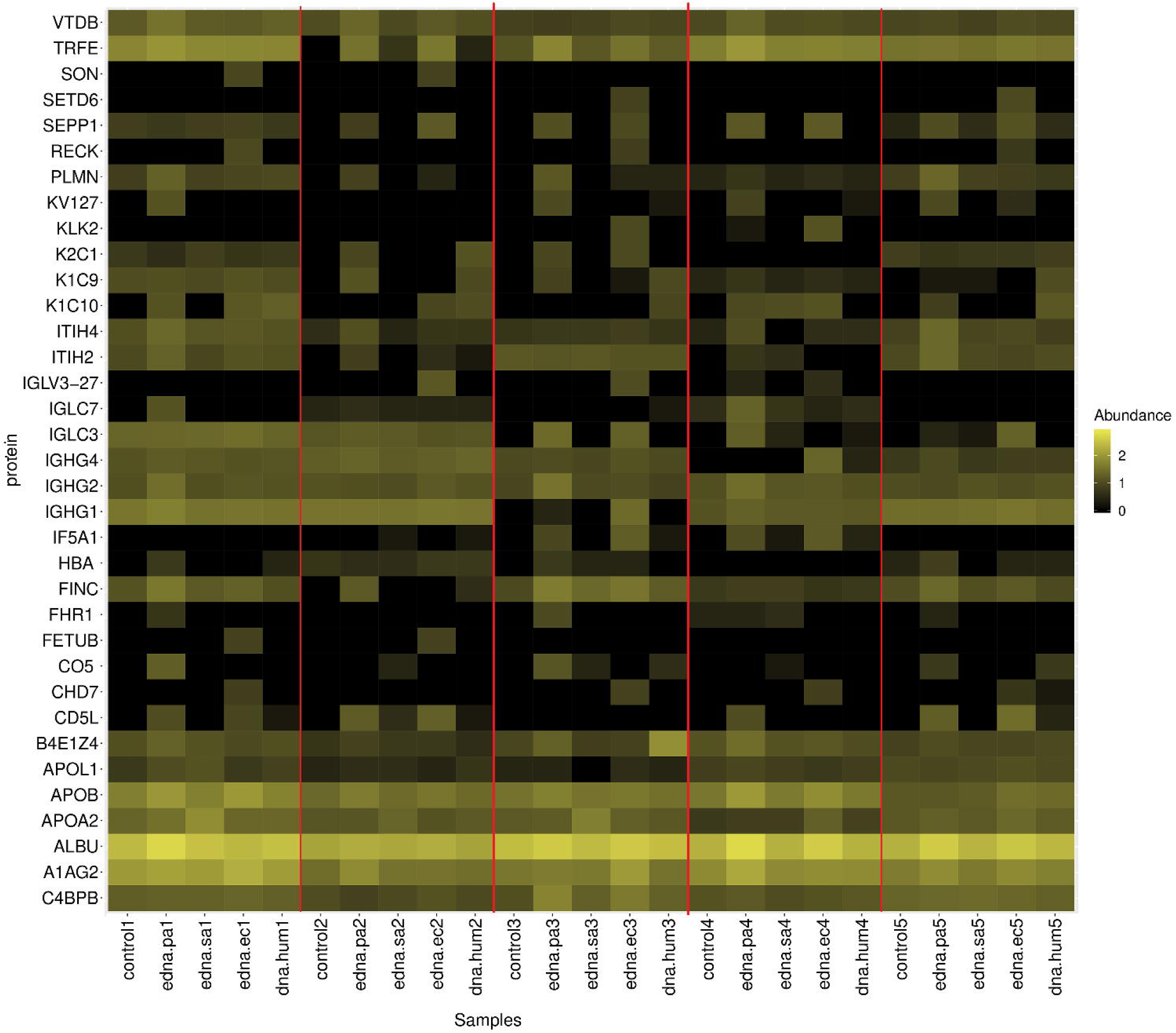
Heatmap of proteins of normal plasma samples that altered their heat resistant characteristics following the treatment with different DNA.

Plasma exposure to the eDNA of *P. aeruginosa* resulted in the formation of 12 heat-resistant proteins, while only some of these proteins, namely K1C10, SEPP1, IGLC3, and IF5A1 acquired heat resistance after treatment with the DNA of another gram-negative bacteria, *E. coli*. The latter, in turn, changed the heat resistance profile of distinct proteins in the same plasma samples. Notably, whereas bacterial eDNA induced heat resistance of a broad spectrum of unrelated proteins, plasma exposure to human DNA only affected the thermal behaviour of a specific group of proteins, i.e., cytoskeletal keratins.

Since prion domains may be responsible for protein heat resistance, we next the employed the prion-prediction PLAAC algorithm to verify the presence of PrDs in proteins exhibiting changes in thermal behaviour following DNA treatment.

We only found PrDs in CHD7 and K1C10, which became heat-resistant following the exposure to *E. coli* eDNA and keratins (K2C1, K1C9, K1C10), which acquired heat resistance upon treatment with both *P. aeruginosa* eDNA and human DNA (Table 3). Notably, these were the only proteins undergoing thermal behaviour alterations following exposure to human DNA.

**Table 3.**
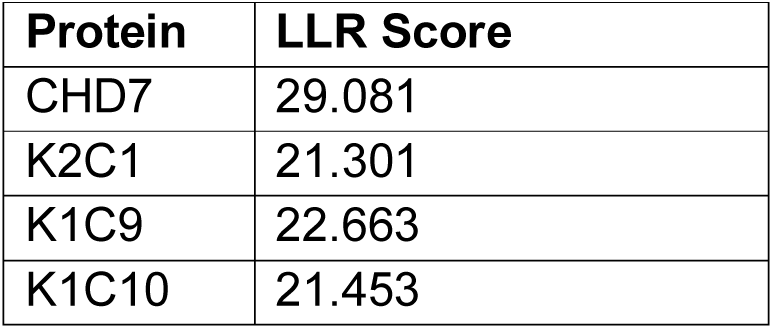
Log-likelihood ratio (LLR) score for PrD predictions in plasma proteins that became heat-resistant following DNA treatment.

| Protein | LLR Score |
| --- | --- |
| CHD7 | 29.081 |
| K2C1 | 21.301 |
| K1C9 | 22.663 |
| K1C10 | 21.453 |

We next analysed the association between DNA-induced changes in protein thermal behaviour and human diseases. Surprisingly, the majority of these proteins had been found associated with cancer progression and some of them are used as a tumour markers (Table 4).

**Table 4.** Association between proteins exhibiting DNA-induced changes in thermal behaviour and human diseases.

| Disease | Proteins | References |
| --- | --- | --- |
| Pancreatic cancer | • Serotransferrin<br>• Complement factor H-related protein<br>• Plasma protease C1 inhibitor<br>• Fibronectin<br>• Immunoglobulin lambda constant 7<br>• C4b-binding protein alpha chain<br>• Selenoprotein P | 29-39 |
| Colorectal cancer | • APOB<br>• SETD6 | 40 – 42 |
|  | • Reversion-inducing cysteine-rich protein with Kazal motifs (RECK) | . |
| Ovarian cancer | • Hemoglobin-α<br>• Eukaryotic translation initiation factor 5A-1<br>• Fibronectin<br>• Inter-α-trypsin inhibitor heavy chain H4 fragment | 43- 46 |
| Breast cancer | • Inter-α-trypsin inhibitor heavy chain H4 fragment | 46 |
| Lung Cancer | • ITIH4<br>• Complement Factor H<br>• Plasma protease C1 inhibitor<br>• Immunoglobulin lambda constant 7<br>• CD5L | 47- 51 |
| hairy cell leukemia. | • Immunoglobulin kappa variable 1-27 | 52 |
| melanoma | • CD5 antigen-like<br>• Keratin, type I cytoskeletal 9 | 53, 54 |
| Prostatic cancer | • Selenoprotein P<br>• Kallikrein 2<br>• apolipoprotein A-II | 55 - 59 |
| Bladder cancer | • SETD6<br>• Complement factor H-related protein | 60,61 |
| Thalassemia | • HBA | 62 |

**Table 4.** Characteristics of subjects and plasma samples.

| Probe | Gender | Age | Tumour Stage | Tumour site | Tumour type |
| --- | --- | --- | --- | --- | --- |
| Control 1 | F | 64 | NA | NA | NA |
| Control 2 | F | 55 | NA | NA | NA |
| Control 3 | M | 57 | NA | NA | NA |
| Control 4 | M | 62 | NA | NA | NA |
| Control 5 | M | 58 | NA | NA | NA |
| Pancreatic cancer 1 | F | 63 | T3N1M1 | Head | Adenocarcinoma |
| Pancreatic cancer 2 | M | 57 | T3N1M1 | Head | Adenocarcinoma |
| Pancreatic cancer 3 | F | 56 | T3N1M1 | Head | Adenocarcinoma |
| Pancreatic cancer 4 | F | 69 | T3N1M1 | Head | Adenocarcinoma |
| Pancreatic cancer 5 | M | 61 | T3N1M1 | Head | Adenocarcinoma |

Intriguingly, some of these cancer-related proteins are also known to be associated with other multifactorial diseases. For example, ITIH4 is associated with schizophrenia and CHD7 is known to be implicated in autism [63-65].

### Comparison of heat-resistant proteome profile in normal, DNA-treated, and pancreatic cancer plasma

We then examined the changes in protein thermal behaviour induced by DNA in normal plasma and compared the resulting pattern with the heat-resistant proteome of patients with pancreatic cancer (Table 4).

After boiling, the plasma samples of patients with pancreatic cancer were characterized for the presence of heat-resistant proteins. Notably, the majority of these proteins were the same that became heat-resistant in normal plasma exposed to DNA treatment. This might suggest that DNA exposure is responsible for cancer-related alterations in the thermal behaviour of specific proteins.

To further explore the relationship between the heat-resistant proteome of patients with pancreatic cancer and the proteome changes induced by DNA in the plasma of healthy individuals, we analysed the scaled spectral counts of the identified heat-resistant proteins of both groups by principal component analysis (PCA) (Figure 2A).

**Figure 2.**
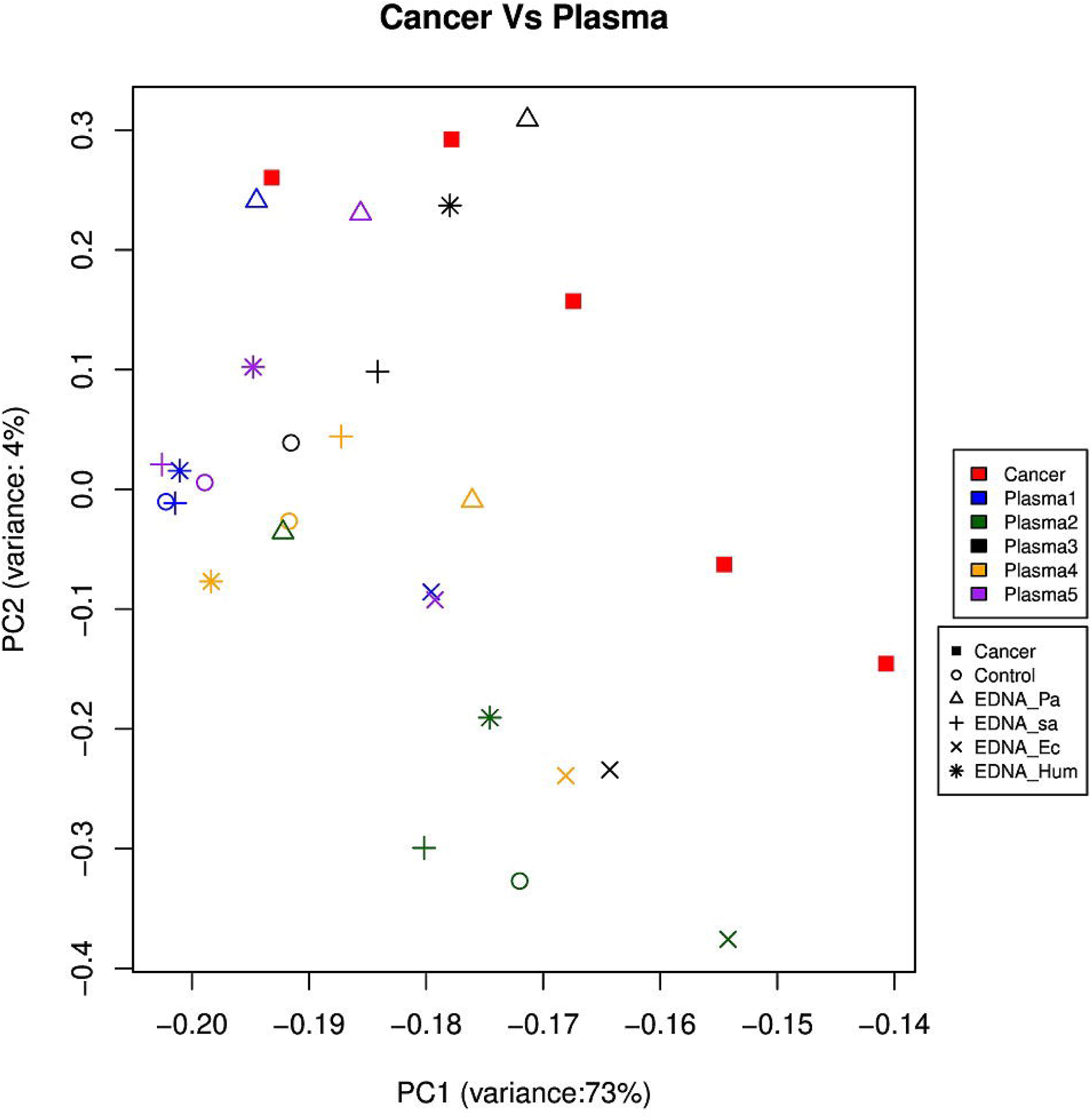
Principal component analysis (PCA) and heat map of proteome data. The heat map represents the relative effects of DNA from different sources on the proportion of heat-resistant proteins in normal plasma. The colour intensity is a function of protein spectrum counts, with bright yellow and black indicating maximal counts and lack of detection, respectively. A) Principal component analysis reflecting the similarities between the heat-resistant proteome of pancreatic cancer plasma and that of plasma from healthy controls following treatment with different DNAs (by LC/MS). The strongest similarities between the plasma of cancer patients and that of healthy subjects after exposure to the eDNA of *P. aeruginosa* are shown B) Heat map showing the mean spectrum counts of heat-resistant proteins in normal plasma samples following DNA treatment, and in the plasma of patients with pancreatic cancer. Black colour and yellow colours represent low and high spectral counts, respectively.

The PCA projection demonstrated that the exposure to bacterial DNA (especially the eDNA of *P. aeruginosa*), induces, in the proteome of normal plasma, changes in thermal behaviour.

A heat map based on the highest spectral counts relative to heat-resistant proteins confirmed that treatment of normal plasma with eDNA of *P. aeruginosa* induced a heat-resistant proteome that had a trend (statistically insignificant) more similar to that of plasma from cancer patients than to untreated plasma (Figure 2B).

## Discussion

This study is the first to demonstrate that bacterial eDNA alters the thermal behaviour of specific proteins in human plasma, leading to an increase in the heat-resistant fraction, as well as to the acquisition of heat resistance by proteins that did not exhibit such property prior to DNA exposure.

For treatment of plasma samples, we used a fixed concentration of bacterial or human DNA, 1 µg/mL, which was selected based on previous studies reporting the presence of similar concentrations of circulating cfDNA in patients with cancer [66, 67].

We discovered that bacterial eDNA or human DNA led to the appearance of different heat-resistant proteins, depending on the DNA source.

Furthermore, we identified a differential effect of eDNA from various gram-positive and gram-negative bacteria on the thermal behaviour of plasma proteins. In fact, we surprisingly found that eDNA from different bacteria interact with distinct plasma proteins (Table 1; Supplementary table 1).

Notably, among the 35 identified proteins with increased heat-resistance following DNA exposure, according to literature data and BindUP tool, only 3 have been previously reported to be able to bind nucleic acids, namely, fibronectin, chromodomain-helicase-DNA-binding protein 7, and SON [68-70].

Heat resistance was previously described only for complement factor H and fibronectin, whereas the other proteins found to contain heat-resistant fragments in this study were not known to possess this property [71-73].

Previous studies have shown that one possible mechanism responsible for the acquisition of heat resistance is the formation of β-structures, which confer increased stability to chemical and physical agents [74-76].

Within this framework, we employed a PLAAC algorithm to verify the presence of PrDs in proteins that were found to acquire heat resistance upon DNA exposure [77]. Of the 35 proteins, only a few were predicted to contain PrDs, i.e., cytoskeletal and microfibrillar keratins I and II, and chromodomain-helicase-DNA-binding protein 7.

Notably, these proteins exhibited a high likelihood ratio (LLR between 21 to 29), and therefore were highly probable to display a prion-like behaviour since the lowest LLR value reported for a known prion-forming protein of budding yeast is ∼21.0 [78].

Interestingly, the PrD-containing K2C1, K1C9, and K1C10 were the only proteins that were found to acquire heat resistance following treatment with human DNA. Another DNA that induced heat resistance in proteins with PrDs was *P. aeruginosa* eDNA. *E. coli* DNA altered heat resistance for K1C10 and has also altered the heat resistance properties of CHD7.

Changes in the thermal behaviour of PrD-containing proteins could be due to the DNA-induced formation of stable β-structures, consistently with the known ability of bacterial DNA in microbial biofilms to interact with bacterial alpha-amyloid, leading to the formation of cross-β structures [79, 80].

However, the majority of proteins undergoing eDNA-dependent changes in heat resistance identified in the current study did not contain PrDs. This suggested that eDNA caused a PrD-independent induction of heat resistance in these proteins. We named proteins undergoing DNA-dependent changes in thermal behaviour, albeit in the absence of prion-like structure, “Tetz-proteins”.

Next, we analysed the association between proteins that acquired DNA-induced heat resistance and human pathologies. According to the literature, many of them were related to a variety of diseases, predominantly cancers.

For example, C4BP, CHD7, and KLK2 are known as tumour biomarkers, while CD5L, EIF5A1, FINC, and SEPP1 are not only described as biomarkers but also known to participate in tumour progression [31, 33, 57, 81].

Our findings suggested a novel role of bacterial eDNA in disease development, and cancer development in particular, consistent with its reported presence in the systemic circulation in association with cancer and other human diseases [20, 82]. Indeed, recent studies have shown that patients with non-infectious early-onset cancer display elevated plasma levels of eDNA from bacteria, particularly *Pseudomonas spp.* and *Pannonibacter* spp. [20].

We next studied the presence and composition of heat-resistant proteins in the plasma of patients with pancreatic cancer. The cancer was selected since the majority of proteins that became heat-resistant following bacterial eDNA exposure had been found to be associated with this pathology. The heat-resistant proteins identified by boiling pancreatic cancer plasma were the same that acquired heat resistance in normal plasma upon exposure to eDNA. We here provide the first demonstration that these proteins, known as cancer markers and associated with tumour progression, possess heat resistance.

Next, the heat-resistant proteome of cancer patients was compared with that of control plasma, before and after the exposure to eDNA. Based on the PCA, after treatment with different DNAs, the heat-resistant proteome of control plasma samples had a statistically insignificant a trend toward increased similarity to that of plasma samples from cancer patients, and this effect was maximal when the eDNA of *P. aeruginosa* was employed. We believe that a larger cohort may provide more statistically significant findings. Notably, CD5L, EIF5A1, FINC, and SEPP1 particularly attracted our attention as, according to the literature, they are associated with tumorigenesis [51, 83-85]. In the present study, heat-resistant fractions of these proteins were identified in pancreatic cancer plasma, but not in normal plasma, and formed only after eDNA treatment, suggesting a role of this conversion in tumorigenesis.

It is tempting to speculate that DNA, including bacterial eDNA, may function as a virulence factor through the interaction with (and the alteration of) Tetz-proteins, including those associated with tumour growth. Therefore, it is possible that under certain conditions, eDNA elevation triggers alterations in plasma proteins that, in turn, may be relevant for tumorigenesis or for other pathologies. Notably, we also found an effect on proteins implicated in neurodegeneration and psychotic disorders. Experiments aimed at the characterization of the pathogenic role of different types of bacterial eDNA are in progress.

## Methods

### Plasma samples

Human plasma samples from 5 healthy donors (age: 57–64 years, 40% females) and 5 patients with clinically diagnosed pancreatic ductal adenocarcinoma (age: 56–69 years, 60% females) were obtained from Bioreclamation IVT (NY, USA) and Discovery Life Sciences (Los Osos, CA). All patients with pancreatic ductal adenocarcinoma had been diagnosed by histological examination and had not undergone surgical treatment, preoperative chemotherapy or radiotherapy. The basic demographic characteristics of the patients are shown in table 4. All samples were obtained with prior informed consent at all facilities. Plasma samples were stored at −80 °C until use.

### Extracellular DNA

Extracellular DNA was extracted from the matrix of *P. aeruginosa* ATCC 27853, *E. coli* ATCC 25922, and *Staphylococcus aureus* ATCC 29213. All bacterial strains were subcultured from freezer stocks onto Columbia agar plates (Oxoid Ltd., London, England) and incubated at 37 °C for 48 h. To extract the extracellular DNA, bacterial cells were separated from the matrix by centrifugation at 5000 g for 10 min at 4 °C. The supernatant was aspirated and filtered through a 0.2-μm-pore-size cellulose acetate filter (Millipore Corporation, USA). eDNA was extracted by using a DNeasy kit (Qiagen), as described by the manufacturer, or by the phenol-chloroform method [86]. Human genomic DNA (Roche Cat#11691112001) was purchased from Sigma (Sigma-Aldrich).

### Plasma exposure to eDNA

DNA was added to plasma samples at the final concentration of 1 µg/mL, incubated at 37 °C for 1 h, and boiled in a water bath at 100 °C for 15 min (by that time all the samples formed clod by coagulated proteins). Samples were cooled at room temperature for 30 min and centrifuged at 5000 g for 10 min at room temperature. The supernatant was aspirated and filtered through a 0.2-μm pore size cellulose acetate filter (Millipore Corporation, USA).

### Protein identification by LS-MS

The filtered protein-containing supernatant was diluted in a final volume of 100 µL using 100 mM ammonium bicarbonate, pH 8, and quantified using a Nanodrop OneC Spectrophotometer (Thermo Fisher Scientific). Cysteine residues were reduced using 5 mM dithiothreitol at room temperature for 1.5 h and alkylated with 10 mM iodoacetamide at room temperature for 45 min in the dark. Proteins were then digested using modified trypsin (Promega, P/N V5113) at a 1:20 (w/w) enzyme:protein ratio for 16 h at 22°C room temperature. After digestion, peptides were acidified to pH 3 with formic acid and desalted using Pierce Peptide Desalting Spin Columns (P/N 89852), according to the manufacturer’s protocol. Eluted, desalted peptides were dried down to completion using a Labconco speedvac concentrator, resuspended in 0.1% formic acid and quantified again using a Nanodrop OneC Spectrophotometer [87]. For sample injection and mass analysis, peptides were diluted to a final concentration of 500 ng/µL using 0.1% formic acid in water to provide a total injection amount of 500 ng in a 1 µL of sample loop. Peptides were separated and their mass analysed using a Dionex UltiMate 3000 RSLCnano ultra-high performance liquid chromatograph (UPLC) coupled to a Thermo Scientific Q Exactive HF hybrid quadrupole-orbitrap mass spectrometer (MS). A 1.5 hr reversed-phase UPLC method was used to separate peptides using a nanoEASE m/z peptide BEH C18 analytical column (Waters, P/N 186008795). The MS method included top 15 data-dependent acquisition for interrogation of peptides by MS/MS using HCD fragmentation. All raw data were searched against the human Uniprot protein database (UP000005640, accessed Apr 22, 2017) using the Andromeda search algorithm within the MaxQuant suite (v 1.6.0.1) [88, 89]. The search results were filtered to a 1 % FPR and visualized using Scaffold (v4, Proteome Software).

A cut-off of at least 5 spectral counts per probe was applied for protein selection [90-92]. The obtained data were used to generate a heatmap. The abundance values were log converted (zero values were replaced with infinitely small number “1”) and plotted with R-statistical computing (https://www.r-project.org/), using the “levelplot” package. The colour key indicates a range between the lowest (black) and the highest (yellow) values. Principal components analysis was performed using the prcomp function with default parameters (zero values were replaced with 1) of the R software (https://www.r-project.org/).

### Identification of prion-like domains (PrDs) in proteins

The presence of prion-like domains in the proteins was assessed using the PLAAC prion prediction algorithm, which establishes the prionogenic nature on the basis of the asparagine (Q) and glutamine (N) content, using the hidden Markov model (HMM) [77, 93]. The output probabilities for the PrD states in PLAAC were estimated based on the amino acid frequencies in the PrDs of *Saccharomyces cerevisiae*. Here, we used Alpha□=□0.0, representing species-independent scanning, to identify the PrDs.

### Data availability

The other sequencing datasets generated and/or analysed during the current study are available from the corresponding author on reasonable request.

## Acknowledgements

All mass spectrometry experiments were conducted by Dr. Jeremy L. Balsbaugh, Director of the Proteomics & Metabolomics Facility, a laboratory which is part of the University of Connecticut’s Center for Open Research Resources & Equipment.

## Author Contributions

VT designed and conducted the experiments. VT and GT analyzed data and wrote the manuscript.

## References

1. Cryan JF, Dinan TG. Mind-altering microorganisms: the impact of the gut microbiota on brain and behaviour. Nature reviews neuroscience. 2012 Oct;13(10):701.

2. Clemente JC, Ursell LK, Parfrey LW, Knight R. The impact of the gut microbiota on human health: an integrative view. Cell. 2012 Mar 16;148(6):1258–70.

3. Tetz, G. et al. Bacteriophages as potential new mammalian pathogens. Sci. Rep. 7, (2017).

4. Tetz G, Brown SM, Hao Y, Tetz V. Type 1 Diabetes: an Association Between Autoimmunity the Dynamics of Gut Amyloid-producing E. coli and Their Phages. BioRxiv. 2018 Jan 1:433110.

5. Tetz, G., Brown, S., Hao, Y. and Tetz, V., 2018. Parkinsons disease and bacteriophages as its overlooked contributors. Scientific reports, 8: 10812.

6. Gasiorowski K, Brokos B, Echeverria V, Barreto GE, Leszek J. RAGE-TLR crosstalk sustains chronic inflammation in neurodegeneration. Molecular neurobiology. 2018 Feb 1;55(2):1463–76.

7. Whitchurch CB, Tolker-Nielsen T, Ragas PC, Mattick JS. Extracellular DNA required for bacterial biofilm formation. Science. 2002 Feb 22;295(5559):1487.

8. Molin S, Tolker-Nielsen T. Gene transfer occurs with enhanced efficiency in biofilms and induces enhanced stabilisation of the biofilm structure. Current opinion in biotechnology. 2003 Jun 1;14(3):255–61.

9. Steinberger RE, Holden PA. Extracellular DNA in single-and multiple-species unsaturated biofilms. Applied and environmental microbiology. 2005 Sep 1;71(9):5404–10.

10. Schwartz K, Ganesan M, Payne DE, Solomon MJ, Boles BR. Extracellular DNA facilitates the formation of functional amyloids in S taphylococcus aureus biofilms. Molecular microbiology. 2016 Jan;99(1):123–34.

11. Evans ML, Chapman MR. Curli biogenesis: order out of disorder. Biochimica et Biophysica Acta (BBA)-Molecular Cell Research. 2014 Aug 1;1843(8):1551–8.

12. Wang X, Chapman MR. Curli provide the template for understanding controlled amyloid propagation. Prion. 2008 Apr 1;2(2):57–60.

13. Galea CA, High AA, Obenauer JC, Mishra A, Park CG, Punta M, Schlessinger A, Ma J, Rost B, Slaughter CA, Kriwacki RW. Large-scale analysis of thermostable, mammalian proteins provides insights into the intrinsically disordered proteome. Journal of proteome research. 2008 Dec 9;8(1):211–26.

14. Narasimhan D, Nance MR, Gao D, Ko MC, Macdonald J, Tamburi P, Yoon D, Landry DM, Woods JH, Zhan CG, Tesmer JJ. Structural analysis of thermostabilizing mutations of cocaine esterase. Protein Engineering, Design & Selection. 2010 Apr 30;23(7):537–47.

15. Wang L, Llorente C, Hartmann P, Yang AM, Chen P, Schnabl B. Methods to determine intestinal permeability and bacterial translocation during liver disease. Journal of immunological methods. 2015 Jun 1;421:44–53.

16. Tetz G, Brown SM, Hao Y, Tetz V. Type 1 Diabetes: an Association Between Autoimmunity the Dynamics of Gut Amyloid-producing E. coli and Their Phages. BioRxiv. 2018 Jan 1:433110.

17. Santis A. Intestinal permeability in non-alcoholic fatty liver disease: the gut-liver axis. Reviews on recent clinical trials. 2014 Sep 1;9(3):141–7.

18. Swarup V, Rajeswari MR. Circulating (cell-free) nucleic acids–A promising, non-invasive tool for early detection of several human diseases. FEBS letters. 2007 Mar 6;581(5):795–9.

19. Snyder MW, Kircher M, Hill AJ, Daza RM, Shendure J. Cell-free DNA comprises an in vivo nucleosome footprint that informs its tissues-of-origin. Cell. 2016 Jan 14;164(1-2):57-68.

20. Huang YF, Chen YJ, Fan TC, Chang NC, Chen YJ, Midha MK, Chen TH, Yang HH, Wang YT, Alice LY, Chiu KP. Analysis of microbial sequences in plasma cell-free DNA for early-onset breast cancer patients and healthy females. BMC medical genomics. 2018 Feb;11(1):16.

21. Wu TL, Zhang D, Chia JH, Tsao KC, Sun CF, Wu JT. Cell-free DNA: measurement in various carcinomas and establishment of normal reference range. Clinica chimica acta. 2002 Jul 1;321(1-2):77–87.

22. Emery DC, Shoemark DK, Batstone TE, Waterfall CM, Coghill JA, Cerajewska TL, Davies M, West NX, Allen SJ. 16S rRNA next generation sequencing analysis shows bacteria in Alzheimer’s post-mortem brain. Frontiers in aging neuroscience. 2017 Jun 20;9:195.

23. Zhao WD, Liu DX, Chen YH. Escherichia coli hijack Caspr1 receptor to invade cerebral vascular and neuronal hosts. Microbial Cell. 2018 Sep 3;5(9):418.

24. Dominy SS, Lynch C, Ermini F, Benedyk M, Marczyk A, Konradi A, Nguyen M, Haditsch U, Raha D, Griffin C, Holsinger LJ. Porphyromonas gingivalis in Alzheimer’s disease brains: Evidence for disease causation and treatment with small-molecule inhibitors. Science Advances. 2019 Jan 1;5(1):eaau3333.

25. Amarasekara R, Jayasekara RW, Senanayake H, Dissanayake VH. Microbiome of the placenta in pre-eclampsia supports the role of bacteria in the multifactorial cause of pre-eclampsia. Journal of Obstetrics and Gynaecology Research. 2015 May 1;41(5):662–9.

26. Mydock-McGrane L, Cusumano Z, Han Z, Binkley J, Kostakioti M, Hannan T, Pinkner JS, Klein R, Kalas V, Crowley J, Rath NP. Antivirulence C-mannosides as antibiotic-sparing, oral therapeutics for urinary tract infections. Journal of medicinal chemistry. 2016 Oct 14;59(20):9390–408.

27. Dikshit N, Bist P, Fenlon SN, Pulloor NK, Chua CE, Scidmore MA, Carlyon JA, Tang BL, Chen SL, Sukumaran B. Intracellular uropathogenic E. coli exploits host Rab35 for iron acquisition and survival within urinary bladder cells. PLoS pathogens. 2015 Aug 6;11(8):e1005083.

28. Dalpke A, Frank J, Peter M, Heeg K. Activation of toll-like receptor 9 by DNA from different bacterial species. Infection and immunity. 2006 Feb 1;74(2):940–6.

29. Takata T, Ishigaki Y, Shimasaki T, Tsuchida H, Motoo Y, Hayashi A, Tomosugi N. Characterization of proteins secreted by pancreatic cancer cells with anticancer drug treatment in vitro. Oncology reports. 2012 Dec 1;28(6):1968–76.

30. Bloomston M, Zhou JX, Rosemurgy AS, Frankel W, Muro-Cacho CA, Yeatman TJ. Fibrinogen ? overexpression in pancreatic cancer identified by large-scale proteomic analysis of serum samples. Cancer research. 2006 Mar 1;66(5):2592– 9.

31. Sogawa K, Takano S, Iida F, Satoh M, Tsuchida S, Kawashima Y, Yoshitomi H, Sanda A, Kodera Y, Takizawa H, Mikata R. Identification of a novel serum biomarker for pancreatic cancer, C4b-binding protein α-chain (C4BPA) by quantitative proteomic analysis using tandem mass tags. British journal of cancer. 2016 Oct;115(8):949.

32. Maehara SI, Tanaka S, Shimada M, Shirabe K, Saito Y, Takahashi K, Maehara Y. Selenoprotein P, as a predictor for evaluating gemcitabine resistance in human pancreatic cancer cells. International journal of cancer. 2004 Nov 1;112(2):184–9.

33. Short SP, Williams CS. Selenoproteins in tumorigenesis and cancer progression. In Advances in cancer research 2017 Jan 1 (Vol. 136, pp. 49-83). Academic Press.

34. Seldon CS, Colbert LE, Hall WA, Fisher SB, Yu DS, Landry JC. Chromodomainhelicase-DNA binding protein 5, 7 and pronecrotic mixed lineage kinase domain–like protein serve as potential prognostic biomarkers in patients with resected pancreatic adenocarcinomas. World journal of gastrointestinal oncology. 2016 Apr 15;8(4):358.

35. Pan S, Chen R, Crispin DA, May D, Stevens T, McIntosh MW, Bronner MP, Ziogas A, Anton-Culver H, Brentnall TA. Protein alterations associated with pancreatic cancer and chronic pancreatitis found in human plasma using global quantitative proteomics profiling. Journal of proteome research. 2011 Mar 28;10(5):2359–76.

36. Crnogorac-Jurcevic T, Missiaglia E, Blaveri E, Gangeswaran R, Jones M, Terris B, Costello E, Neoptolemos JP, Lemoine NR. Molecular alterations in pancreatic carcinoma: expression profiling shows that dysregulated expression of S100 genes is highly prevalent. The Journal of Pathology: A Journal of the Pathological Society of Great Britain and Ireland. 2003 Sep;201(1):63–74.

37. Zhao J, Simeone DM, Heidt D, Anderson MA, Lubman DM. Comparative serum glycoproteomics using lectin selected sialic acid glycoproteins with mass spectrometric analysis: application to pancreatic cancer serum. Journal of proteome research. 2006 Jul 7;5(7):1792–802.

38. Nie S, Yin H, Tan Z, Anderson MA, Ruffin MT, Simeone DM, Lubman DM. Quantitative analysis of single amino acid variant peptides associated with pancreatic cancer in serum by an isobaric labeling quantitative method. Journal of proteome research. 2014 Nov 24;13(12):6058–66.

39. Cecconi D, Palmieri M, Donadelli M. Proteomics in pancreatic cancer research. Proteomics. 2011 Feb;11(4):816–28.

40. Martín-Morales L, Feldman M, Vershinin Z, Garre P, Caldés T, Levy D. SETD6 dominant negative mutation in familial colorectal cancer type X. Human molecular genetics. 2017 Aug 30;26(22):4481–93.

41. Oshima, T., Kunisaki, C., Yoshihara, K., Yamada, R., Yamamoto, N., Sato, T., Makino, H., Yamagishi, S., Nagano, Y., Fujii, S. and Shiozawa, M., 2008. Clinicopathological significance of the gene expression of matrix metalloproteinases and reversion-inducing cysteine-rich protein with Kazal motifs in patients with colorectal cancer: MMP-2 gene expression is a useful predictor of liver metastasis from colorectal cancer. Oncology reports, 19(5), pp.1285–1291.

42. Borgquist S, Butt T, Almgren P, Shiffman D, Stocks T, Orho-Melander M, Manjer J, Melander O. Apolipoproteins, lipids and risk of cancer. International journal of cancer. 2016 Jun 1;138(11):2648–56.

43. Woong-Shick A, Sung-Pil P, Su-Mi B, Joon-Mo L, Sung-Eun N, Gye-Hyun N, Young-Lae C, Ho-Sun C, Heung-Jae J, Chong-Kook K, Young-Wan K. Identification of hemoglobin-a and-ß subunits as potential serum biomarkers for the diagnosis and prognosis of ovarian cancer. Cancer science. 2005 Mar;96(3):197–201.

44. Zhang, J., Li, X., Liu, X., Tian, F., Zeng, W., Xi, X., & Lin, Y. (2018). EIF5A1 promotes epithelial ovarian cancer proliferation and progression. Biomedicine & Pharmacotherapy, 100, 168–175.

45. Wang JP, Hielscher A. Fibronectin: how its aberrant expression in tumors may improve therapeutic targeting. Journal of Cancer. 2017;8(4):674.

46. Mohamed E, Abdul-Rahman PS, Doustjalali SR, Chen Y, Lim BK, Omar SZ, Bustam AZ, Singh VA, Mohd-Taib NA, Yip CH, Hashim OH. Lectin-based electrophoretic analysis of the expression of the 35 kDa inter-α-trypsin inhibitor heavy chain H4 fragment in sera of patients with five different malignancies. Electrophoresis. 2008 Jun;29(12):2645–50.

47. Heo SH, Lee SJ, Ryoo HM, Park JY, Cho JY. Identification of putative serum glycoprotein biomarkers for human lung adenocarcinoma by multilectin affinity chromatography and LC-MS/MS. Proteomics. 2007 Dec;7(23):4292–302.

48. Ajona D, Castano Z, Garayoa M, Zudaire E, Pajares MJ, Martinez A, Cuttitta F, Montuenga LM, Pio R. Expression of complement factor H by lung cancer cells: effects on the activation of the alternative pathway of complement. Cancer Research. 2004 Sep 1;64(17):6310–8.

49. Sun Y, Liu S, Qiao Z, Shang Z, Xia Z, Niu X, Qian L, Zhang Y, Fan L, Cao CX, Xiao H. Systematic comparison of exosomal proteomes from human saliva and serum for the detection of lung cancer. Analytica chimica acta. 2017 Aug 22;982:84–95.

50. Zeng X, Hood BL, Zhao T, Conrads TP, Sun M, Gopalakrishnan V, Grover H, Day RS, Weissfeld JL, Wilson DO, Siegfried JM. Lung cancer serum biomarker discovery using label-free liquid chromatography-tandem mass spectrometry. Journal of Thoracic Oncology. 2011 Apr 1;6(4):725–34.

51. Li Y, Qu P, Wu L, Li B, Du H et al. (2011) Api6/AIM/Spα/CD5L Overexpression in alveolar type II epithelial cells induces spontaneous lung adenocarcinoma. Cancer Res 71(16):5488–5499

52. Forconi F, Sozzi E, Rossi D, Sahota SS, Amato T, Raspadori D, Trentin L, Leoncini L, Gaidano G, Lauria F. Selective influences in the expressed immunoglobulin heavy and light chain gene repertoire in hairy cell leukemia. haematologica. 2008 May 1;93(5):697–705.

53. Darling VR, Hauke RJ, Tarantolo S, Agrawal DK. Immunological effects and therapeutic role of C5a in cancer. Expert review of clinical immunology. 2015 Feb 1;11(2):255–63.

54. Chen N, Gong J, Chen X, Xu M, Huang Y, Wang L, Geng N, Zhou Q. Cytokeratin expression in malignant melanoma: potential application of in-situ hybridization analysis of mRNA. Melanoma research. 2009 Apr 1;19(2):87–93.

55. Cooper ML, Adami HO, Grönberg H, Wiklund F, Green FR, Rayman MP. Interaction between single nucleotide polymorphisms in selenoprotein P and mitochondrial superoxide dismutase determines prostate cancer risk. Cancer research. 2008 Dec 15;68(24):10171–7.

56. Persson-Moschos ME, Stavenow L, Åkesson B, Lindgärde F. Selenoprotein P in plasma in relation to cancer morbidity in middle-aged Swedish men. Nutrition and cancer. 2000 Jan 1;36(1):19–26.

57. Guerrico AG, Hillman D, Karnes J, Davis B, Gaston S, Klee G. Roles of kallikrein-2 biomarkers (free-hK2 and pro-hK2) for predicting prostate cancer progression-free survival. Journal of circulating biomarkers. 2017 Jul 19;6:1849454417720151.

58. Darson MF, Pacelli A, Roche P, Rittenhouse HG, Wolfert RL, Young CY, Klee GG, Tindall DJ, Bostwick DG. Human glandular kallikrein 2 (hK2) expression in prostatic intraepithelial neoplasia and adenocarcinoma: a novel prostate cancer marker. Urology. 1997 Jun 1;49(6):857–62.

59. Malik G, Ward MD, Gupta SK, Trosset MW, Grizzle WE, Adam BL, Diaz JI, Semmes OJ. Serum levels of an isoform of apolipoprotein A-II as a potential marker for prostate cancer. Clinical Cancer Research. 2005 Feb 1;11(3):1073– 85.

60. Malik G, Ward MD, Gupta SK, Trosset MW, Grizzle WE, Adam BL, Diaz JI, Semmes OJ. Serum levels of an isoform of apolipoprotein A-II as a potential marker for prostate cancer. Clinical Cancer Research. 2005 Feb 1;11(3):1073– 85.

61. Raitanen MP, Marttila T, Nurmi M, Ala-opas M, Nieminen P, Aine R, Tammela TL, Finnbladder Group. Human complement factor H related protein test for monitoring bladder cancer. The Journal of urology. 2001 Feb 1;165(2):374–7.

62. Origa, Raffaella, and Paolo Moi. “Alpha-thalassemia.” (2016).

63. Crawley JN, Heyer WD, LaSalle JM. Autism and cancer share risk genes, pathways, and drug targets. Trends in Genetics. 2016 Mar 1;32(3):139–46.

64. Cooper JD, Han SY, Tomasik J, Ozcan S, Rustogi N, van Beveren NJ, Leweke FM, Bahn S. Multimodel inference for biomarker development: an application to schizophrenia. Translational Psychiatry. 2019 Feb 11;9(1):83.

65. La YJ, Wan CL, Zhu H, Yang YF, Chen YS, Pan YX, Feng GY, He L. Decreased levels of apolipoprotein AI in plasma of schizophrenic patients. Journal of neural transmission. 2007 May 1;114(5):657–63.

66. Wu TL, Zhang D, Chia JH, Tsao KC, Sun CF, Wu JT. Cell-free DNA: measurement in various carcinomas and establishment of normal reference range. Clinica chimica acta. 2002 Jul 1;321(1-2):77–87.

67. Tomochika, S., Iizuka, N., Watanabe, Y., Tsutsui, M., Takeda, S., Yoshino, S., Ichihara, K. and Oka, M., 2010. Increased serum cell-free DNA levels in relation to inflammation are predictive of distant metastasis of esophageal squamous cell carcinoma. Experimental and therapeutic medicine, 1(1), pp.89–92.

68. Khan IM, Fisher RA, Johnson KJ, Bailey ME, Siciliano MJ, Kessling AM, Farrer M, Carritt B, Kamalati T, Buluwela L. The SON gene encodes a conserved DNA binding protein mapping to human chromosome 21. Annals of human genetics. 1994 Jan;58(1):25–34.

69. Paz I, Kligun E, Bengad B, Mandel-Gutfreund Y. BindUP: a web server for non-homology-based prediction of DNA and RNA binding proteins. Nucleic acids research. 2016 May 19;44(W1):W568-74.

70. Si J, Zhao R, Wu R. An overview of the prediction of protein DNA-binding sites. International journal of molecular sciences. 2015 Mar;16(3):5194–215.

71. Kask L, Villoutreix BO, Steen M, Ramesh B, Dahlbäck B, Blom AM. Structural stability and heat-induced conformational change of two complement inhibitors: C4b-binding protein and factor H. Protein Science. 2004 May;13(5):1356–64.

72. Privalov PL. Microcalorimetry of macromolecules: protein folding, multidomain proteins.

73. Kopp A, Hebecker M, Svobodová E, Józsi M. Factor h: a complement regulator in health and disease, and a mediator of cellular interactions. Biomolecules. 2012 Mar;2(1):46–75.

74. Saverioni D, Notari S, Capellari S, Poggiolini I, Giese A, Kretzschmar HA, Parchi P. Analyses of protease resistance and aggregation state of abnormal prion protein across the spectrum of human prions. Journal of Biological Chemistry. 2013 Sep 27;288(39):27972–85.

75. Abd-Elhadi S, Honig A, Simhi-Haham D, Schechter M, Linetsky E, Ben-Hur T, Sharon R. Total and proteinase K-resistant α-synuclein levels in erythrocytes, determined by their ability to bind phospholipids, associate with Parkinson’s disease. Scientific reports. 2015 Jun 11;5:11120.

76. Zheng Z, Zhang M, Wang Y, Ma R, Guo C, Feng L, Wu J, Yao H, Lin D. Structural basis for the complete resistance of the human prion protein mutant G127V to prion disease. Scientific reports. 2018 Sep 4;8(1):13211.

77. Lancaster AK, Nutter-Upham A, Lindquist S, King OD. PLAAC: a web and command-line application to identify proteins with prion-like amino acid composition. Bioinformatics. 2014 May 13;30(17):2501–2.

78. An L, Fitzpatrick D, Harrison PM. Emergence and evolution of yeast prion and prion-like proteins. BMC evolutionary biology. 2016 Dec;16(1):24.

79. Schwartz K, Ganesan M, Payne DE, Solomon MJ, Boles BR. Extracellular DNA facilitates the formation of functional amyloids in S taphylococcus aureus biofilms. Molecular microbiology. 2016 Jan;99(1):123–34.

80. Gallo PM, Rapsinski GJ, Wilson RP, Oppong GO, Sriram U, Goulian M, Buttaro B, Caricchio R, Gallucci S, Tükel Ç. Amyloid-DNA composites of bacterial biofilms stimulate autoimmunity. Immunity. 2015 Jun 16;42(6):1171–84.

81. Sanjurjo L, Aran G, Téllez É, Amézaga N, Armengol C, López D, Prats C, Sarrias MR. CD5L promotes M2 macrophage polarization through autophagy-mediated upregulation of ID3. Frontiers in immunology. 2018 Mar 12;9:480.

82. Meisel M, Hinterleitner R, Pacis A, Chen L, Earley ZM, Mayassi T, Pierre JF, Ernest JD, Galipeau HJ, Thuille N, Bouziat R. Microbial signals drive pre-leukaemic myeloproliferation in a Tet2-deficient host. Nature. 2018 May;557(7706):580.

83. Mathews MB, Hershey JW. The translation factor eIF5A and human cancer. Biochimica et Biophysica Acta (BBA)-Gene Regulatory Mechanisms. 2015 Jul 1;1849(7):836–44.

84. Han SW, Roman J. Fibronectin induces cell proliferation and inhibits apoptosis in human bronchial epithelial cells: pro-oncogenic effects mediated by PI3-kinase and NF-κB. Oncogene. 2006 Jul;25(31):4341.

85. Barrett CW, Short SP, Williams CS. Selenoproteins and oxidative stress-induced inflammatory tumorigenesis in the gut. Cellular and molecular life sciences. 2017 Feb 1;74(4):607–16.

86. Maniatis T. Fritsch, EF and Sambrook, J.: Molecular Cloning. A Laboratory Manual. New York. Cold Spring Harbor Laboratory, Cold Spring Harbor. 1982.

87. Dayon L, Kussmann M. Proteomics of human plasma: A critical comparison of analytical workflows in terms of effort, throughput and outcome. EuPA Open Proteomics. 2013 Jan 1;1:8–16.

88. Cox J, Neuhauser N, Michalski A, Scheltema RA, Olsen JV, Mann M. Andromeda: a peptide search engine integrated into the MaxQuant environment. Journal of proteome research. 2011 Feb 22;10(4):1794–805.

89. “UniProt: the universal protein knowledgebase.” Nucleic acids research 45, no. D1 (2016): D158–D169.

90. Sigdel TK, Kaushal A, Gritsenko M, Norbeck AD, Qian WJ, Xiao W, Camp DG, Smith RD, Sarwal MM. Shotgun proteomics identifies proteins specific for acute renal transplant rejection. PROTEOMICS–Clinical Applications. 2010 Jan;4(1):32–47.

91. Tu C, Rudnick PA, Martinez MY, Cheek KL, Stein SE, Slebos RJ, Liebler DC. Depletion of abundant plasma proteins and limitations of plasma proteomics. Journal of proteome research. 2010 Aug 31;9(10):4982–91.

92. Li M, Gray W, Zhang H, Chung CH, Billheimer D, Yarbrough WG, Liebler DC, Shyr Y, Slebos RJ. Comparative shotgun proteomics using spectral count data and quasi-likelihood modeling. Journal of proteome research. 2010 Jul 19;9(8):4295–305.

93. Eddy, S. R. (1998). Profile hidden Markov models. Bioinformatics 14, 755–763. doi: 10.1093/bioinformatics/14.9.755

